# Population-specific sequence and expression differentiation in Europeans

**DOI:** 10.1101/749499

**Authors:** Xueyuan Jiang, Raquel Assis

## Abstract

Much of the enormous phenotypic variation observed across human populations is thought to have arisen from events experienced as our ancestors peopled different regions of the world. However, little is known about the genes involved in these population-specific adaptations. Here we explore this problem by simultaneously examining population-specific sequence and expression differentiation in four human populations. In particular, we design a branch-based statistic to estimate population-specific differentiation in four populations, and apply this statistic to single nucleotide polymorphism (SNP) and RNA-seq data from Italian, British, Finish, and Yoruban populations. As expected, genome-wide estimates of sequence and expression differentiation each independently recapitulate the known demographic history of these four human populations, highlighting the utility of our statistic for identifying genic targets of population-specific adaptations. Application of our statistic reveals that genes containing large copy number variations (CNVs) have elevated levels of population-specific sequence and expression differentiation, consistent with the hypothesis that gene turnover is a key reservoir of adaptive variation. Further, European genes displaying population-specific sequence and expression differentiation are enriched for functions related to epigenetic regulation, immunity, and reproduction. Together, our findings illustrate that population-specific sequence and expression differentiation in humans may preferentially target genes with CNVs and play important roles in a diversity of adaptive and disease-related phenotypes.

## 1. Introduction

Human phenotypes vary widely across the globe. In particular, geographically separated populations often differ in skin pigmentation (Loomis 1967), hair color (Rees 2003), tooth morphology (Scott and Turner 1997; Hanihara and Ishida 2005), surface area to body mass ratio (Katzmarzyk and Leonard 1998), and predisposition to diseases (Frank 2004). Much of this phenotypic variation is thought to have arisen due to a diversity of selective pressures experienced as early humans peopled the world and encountered novel environments (Sabeti et al. 2002; Voight et al. 2006), food sources (Sabeti et al. 2002), and pathogens (Diamond 2002; Jobling et al. 2013). As a result, uncovering the genetic targets of phenotypic variation among human populations is critical both for understanding past human adaptations (Sabeti et al. 2002) and for advancing future biomedical research (Jorde et al. 2001; Akey et al. 2004).

Due to the abundance of whole-genome sequence and polymorphism data for many human populations (Cann et al. 2002; International HapMap 3 Consortium 2010; The 1000 Genome Projects Consortium 2015), much work in the past several years has focused on elucidating and understanding sequence differentiation that occurred during human evolution (Li et al. 2008; Pickrell et al. 2009; Field et al. 2016). A common summary statistic for estimating sequence distances between two populations is the fixation index, *F*_ST_ (Wright 1951), which has been used to infer human demographic history (Hinds et al. 2005; Holsinger and Weir 2009; Keinan et al. 2009; Patterson et al. 2012; The 1000 Genome Projects Consortium 2015) and identify loci that may be targets of natural selection (Bowcock et al. 1991; Akey et al. 2002; Bersaglieri et al. 2004). However, because *F*_ST_ is a pairwise metric, it cannot identify the directionality of sequence differentiation nor be used as sole evidence for natural selection (Yi et al. 2010). To address this issue, Yi et al. (Yi et al. 2010) developed the Population Branch Statistic (PBS), a summary statistic that utilizes pairwise *F*_ST_ values among three populations to quantify sequence differentiation along each branch of their corresponding three-population tree. Genes with large PBS values on one branch represent loci that underwent population-specific sequence differentiation consistent with positive selection (Yi et al. 2010). PBS has been applied to corroborate previously established targets of selection, including genes associated with skin pigmentation (Lamason et al. 2005) and dietary fat sources (Mathias et al. 2012), as well as to identify novel candidates for high-altitude adaptation in Tibetans (Yi et al. 2010).

However, because natural selection acts on phenotypes, analysis of sequence data only enables assessment of its indirect effects. For this reason, it may be advantageous to study selection more directly by exploiting the recent availability of RNA-seq data for several human populations (Lappalainen et al. 2013). Specifically, phenotypic evolution is thought to often occur through modifications in gene expression (King and Wilson 1975; Wang et al. 1996; Wray et al. 2003; Carroll 2005; Carroll 2008; Raj et al. 2010). Thus, studying gene expression differentiation among human populations may increase power for identifying loci underlying population-specific phenotypic variation. Indeed, like genetic differentiation, gene expression levels vary considerably across human populations (Cheung et al. 2005; Stranger et al. 2007) and often reflect population structure (Brown et al. 2018). Moreover, human genes with large PBS values are enriched for expression quantitative trait loci (eQTLs; Quiver and Lachance 2018).

In the present study, we simultaneously explore population-specific sequence and expression differentiation in four human populations: the Toscani in Italia (TSI), British in England and Scotland (GBR), Finnish in Finland (FIN), and Yoruba in Nigeria (YRI). For these analyses, we employ single nucleotide polymorphism (SNP; The 1000 Genome Projects Consortium 2015) and RNA-seq (Lappalainen et al. 2013) data from each population. First, we use *F*_ST_ (Wright 1951) and its analogue for estimating quantitative trait differentiation, *P*_ST_ (Leinonen et al. 2006), to quantify and examine genome-wide patterns of sequence and expression differentiation in the four human populations. Next, we adapt the approach of PBS (Yi et al. 2010) to *P*_ST_, as well as extend its computation to a four-population tree, enabling us to estimate both sequence and expression differentiation in each of the four human populations. Last, we apply these branch-based statistics to study population-specific sequence and expression differentiation in Europeans and uncover candidate genes and functional modules underlying adaptation in TSI, GBR, and FIN populations.

## 2. Results

### 2.1 Genome-wide patterns of sequence and expression differentiation in four human populations

A first goal of our study was to estimate sequence and expression differentiation among TSI, GBR, FIN, and YRI populations. To address this problem, we used SNP data (The 1000 Genome Projects Consortium 2015) to calculate the *F*_ST_ (Wright 1951), and RNA-seq data (Lappalainen et al. 2013) to calculate the *P*_ST_ (Leinonen et al. 2006), of every gene between each pair of the four human populations. We calculated *F*_ST_ using Hudson’s formula (Hudson et al. 1992) and computed the ratio of averages to minimize bias (Reynolds et al. 1983; Weir and Cockerham 1984; International HapMap 3 Consortium 2010; Bhatia et al. 2013; see Materials and Methods for details). Due to environmental effects on *P*_ST_, we followed the approach of Leinonen et al. (2006) in calculating *P*_ST_ under two contrasting scenarios: one in which environmental and non-additive genetic effects account for half of the observed expression variation (*h*^*2*^ = 0.5), and a second in which only additive genetic effects contribute to the observed expression variation (*h*^*2*^ = 1; see Materials and Methods for details). Examinations of Pearson’s linear (*r*) and Spearman’s nonlinear (*ρ*) correlations revealed small but significantly positive relationships between *F*_ST_ and *P*_ST_ in TSI-FIN, TSI-YRI, GBR-YRI, and FIN-YRI population pairs (Tables S1-S2), consistent with previous observations that sequence and expression differentiation are weakly or moderately associated (Makova and Li 2003; Nuzhdin et al. 2004; Sartor et al. 2006; Hunt et al. 2012; Assis and Bachtrog 2013; Assis and Bachtrog 2015).

To explore genome-wide patterns of sequence and expression differentiation among the four human populations, we independently used *F*_ST_ and *P*_ST_ to construct gene trees and then infer population trees supported by majorities of these gene trees (see Materials and Methods for details). Population trees obtained from *F*_ST_ and *P*_ST_ (*h*^*2*^ = 0.5 and *h*^*2*^ =1) have the same topology (Figure 1), indicating that there is consistency between relationships inferred from genome-wide patterns of sequence and expression differentiation despite their weak correlations with one another. Moreover, as expected due to increased noise in gene expression data (Raser and O’Shea 2005), gene trees obtained from *P*_ST_ (*h*^*2*^ = 0.5 and *h*^*2*^ =1) are more variable than those derived from *F*_ST_. Further, the topology of the population trees recapitulates human demographic history, in that TSI and GBR populations are most closely related to one another, the FIN population is an outgroup to TSI and GBR, and the YRI population is an outgroup to all three European populations. These results mirror those from similar studies of *F*_ST_ (Hinds et al. 2005; Holsinger and Weir 2009; Keinan et al. 2009; Patterson et al. 2012; The 1000 Genome Projects Consortium 2015), as well as findings that gene expression data often display population structure comparable to that of sequence data (Cheung et al. 2005; Stranger et al. 2007; Brown et al. 2018).

**Figure 1.**
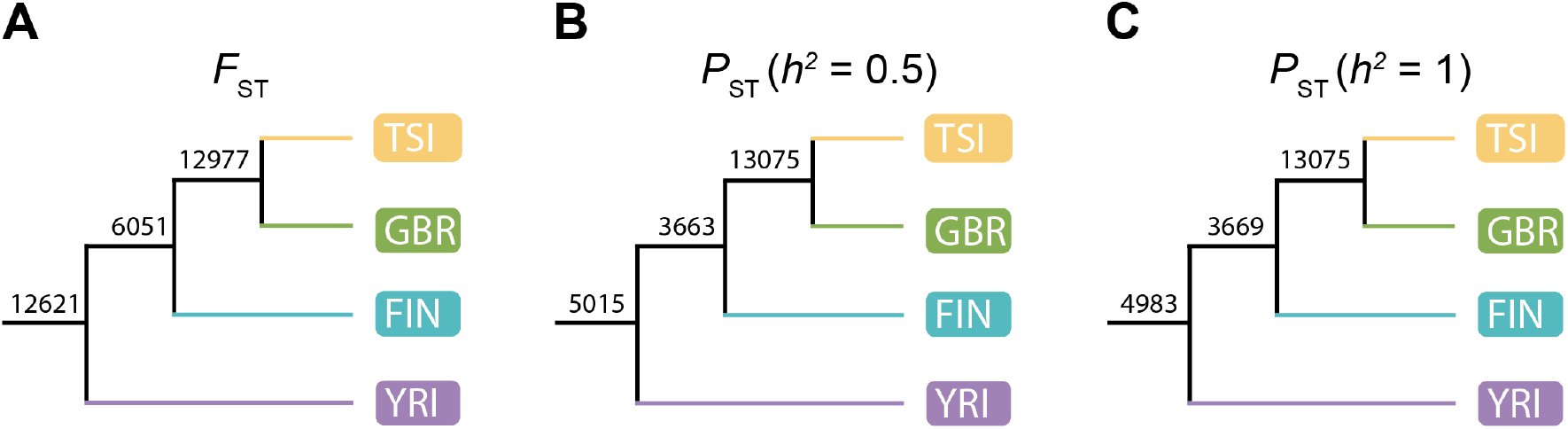
Relationships among TSI, GBR, FIN, and YRI populations inferred from genome-wide patterns of sequence and expression differentiation. Population trees supported by the majority of gene trees built using (*A*) *F*_ST_, (*B*) *P*_ST_ with *h*^*2*^ = 0.5, and (*C*) *P*_ST_ with *h*^*2*^ = 1. Numbers indicate occurrences of corresponding nodes in all gene trees (see Materials and Methods for details).

### 2.2 Estimation of population-specific sequence and expression differentiation on a four-population tree

Next, we sought to quantify population-specific sequence and expression differentiation of genes in each of the four human populations. For a three-population tree, population-specific sequence differentiation of a gene along each branch can be estimated with PBS (Yi et al. 2010) (Figure 2A), which applies Equation 11.20 in Felsenstein (Felenstein 2004) to *F*_ST_. In particular, considering the unrooted three-population tree shown in Figure 2A, the PBS value of a particular gene in population W is estimated as 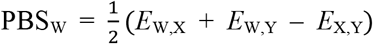, where *E*_W,X_, *E*_W,Y_, and *E*_X,Y_ denote log-transformed *F*_ST_ between populations W and X, W and Y, and X and Y, respectively (Yi et al. 2010; see Materials and Methods for details). In a recent study, Equation 11.20 in Felsenstein (Felenstein 2004) was also applied to expression distances between orthologous genes to estimate branch lengths corresponding to lineage-specific expression divergence on a three-species tree (Assis 2018). Analogously, by substituting *P*_ST_ for *F*_ST_ in the formula for PBS (Yi et al. 2010), we can obtain the PBS corresponding to gene expression differentiation in population W on the three-population tree. To distinguish between these two PBS in our study, we will refer to the calculation with *F*_ST_ as the “sequence PBS”, and the calculation with *P*_ST_ as the “expression PBS”.

**Figure 2.**
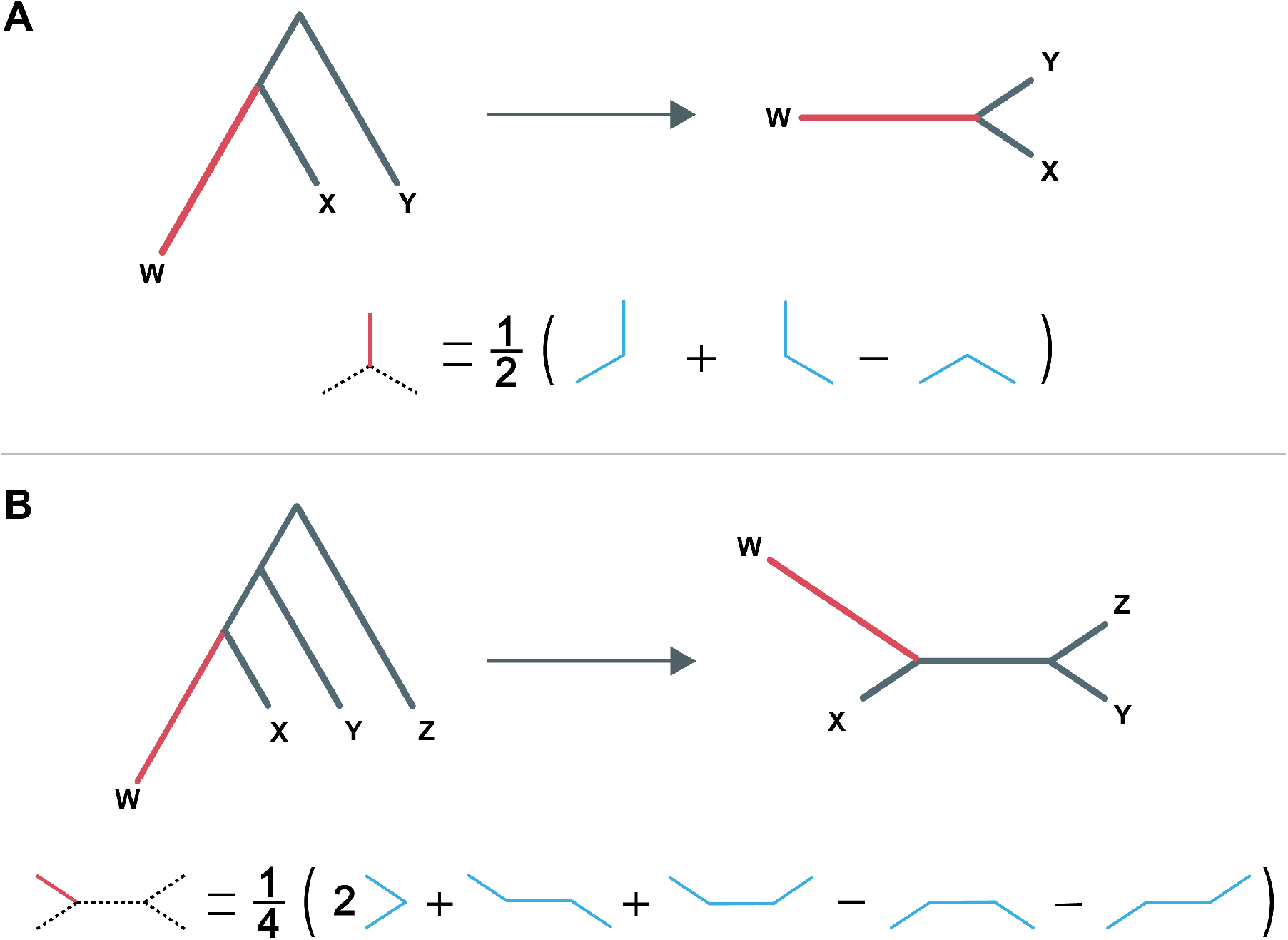
Schematic of PBS calculations. (*A*) Application of PBS_3_ to population W in a tree relating populations W, X, and Y. The rooted three-population tree is unrooted (top), and the length of branch W is computed by applying the formula shown (bottom) to pairwise sequence (*F*_ST_) or expression (*P*_ST_) distances between populations. (*B*) Application of PBS_4_ to population W in a tree relating populations W, X, Y, and Z. The rooted four-population tree is unrooted (top), and the length of branch W is computed by applying the formula shown (bottom) to pairwise sequence (*F*_ST_) or expression (*P*_ST_) distances between populations.

To enable quantification of population-specific sequence and expression differentiation in four human populations, we further extended the computation of each PBS to a four-population tree (Figure 2B) via application of Equation 12.6 in Felsenstein (Felenstein 2004). Henceforth, we will denote PBS as PBS_3_ when applied to a three-population tree, and as PBS_4_ when applied to a four-population tree. In particular, considering the unrooted four-population tree depicted in Figure 2B, the PBS value of a particular gene in population A can be estimated as 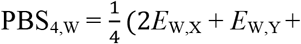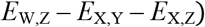, where *E*_W,X_, *E*_W,Y_, *E*_X,Y_, and *E*_X,Z_ denote log-transformed *F*_ST_ or *P*_ST_ of the gene between populations W and X, W and Y, X and Y, and X and Z, respectively (see Materials and Methods for details). We used this formula to compute the sequence PBS_4_ and expression PBS_4_ score of each gene in TSI, GBR, FIN, and YRI populations (Tables S3-S5).

### 2.3 Population-specific sequence and expression differentiation of genes with copy number variations

Gene duplications and deletions are key contributors to human genetic diversity (Sudmant et al. 2015). Moreover, because they are large-scale mutation events that may impact gene dosage, duplications and deletions have been implicated in numerous human diseases (Sebat et al. 2004; Kumar et al. 2008; Sharp et al. 2008; Weiss et al. 2008), as well as in adaptive events in many diverse species (Kaessmann 2010; Chen et al. 2013). As a result, genes containing copy number variations (CNVs) are thought to be more frequently targeted by natural selection than those without CNVs (Freeman et al. 2006; Nguyen et al. 2006). Indeed, genes with CNVs often display signatures of adaptation (Sudmant et al. 2015), and fixation of duplications and deletions has been associated with natural selection in many species (Freeman et al. 2006; Nguyen et al. 2006; Han et al. 2009b; Jiang and Assis 2017). Therefore, we hypothesized that genes with CNVs would have larger sequence and expression PBS_4_ values than genes without CNVs.

To test this hypothesis, we compared the distributions of maximum PBS_4_ values of genes with and without known human CNVs larger than 50bp (Figure 3; MacDonald et al. 2013; see Materials and Methods for details). Consistent with our prediction, both sequence and expression PBS_4_ values are significantly elevated in genes with CNVs (Figure 3; *P* < 0.05 for all pairwise comparisons, two-sample permutation tests; see Materials and Methods for details). Therefore, genes containing large CNVs appear to undergo increased population-specific sequence and expression differentiation, supporting the hypothesis that both their sequences and expression patterns may be targeted by natural selection.

**Figure 3.**
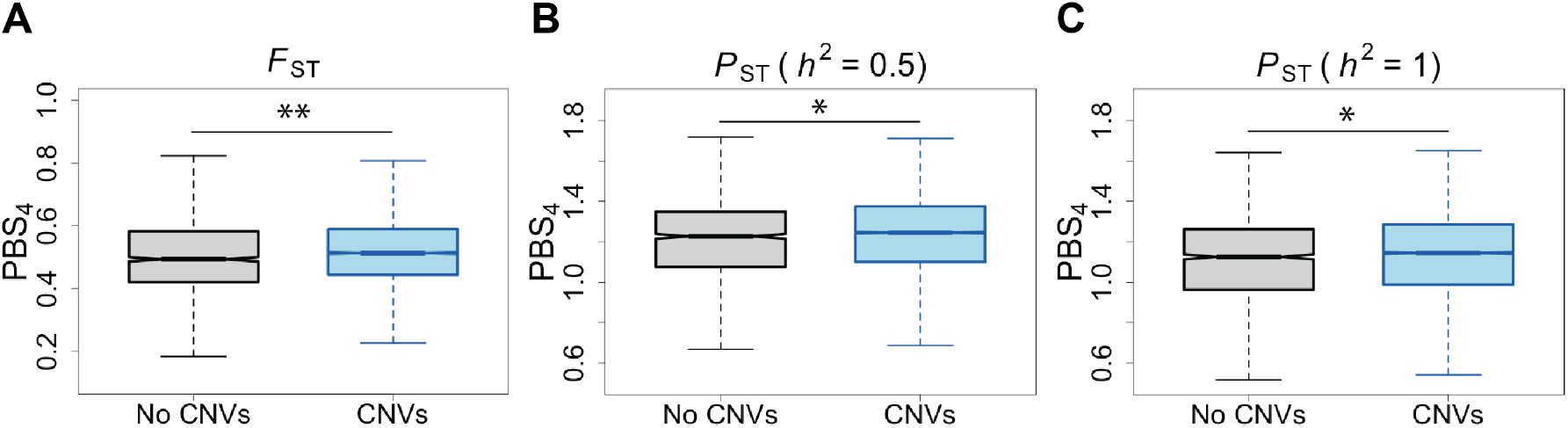
PBS_4_ values of genes with and without CNVs. Distributions of cube root transformed (*A*) sequence PBS_4_ values, (*B*) expression PBS_4_ values with *h*^*2*^ = 0.5, and (*C*) expression PBS_4_ values with *h*^*2*^ = 1 of genes with (blue) and without (gray) CNVs. * *P* < 0.05, ** *P* < 0.001. (See Materials and Methods for details).

### 2.4 Relationships of population-specific sequence and expression differentiation to gene function in Europeans

A natural question that emerges from this analysis is what types of functional modules underlie population-specific sequence and expression differentiation in human populations. In addressing this question, it was important to exclude YRI, as it is an outgroup to the three European populations and therefore contains greater overall population-specific sequence and expression differentiation that is difficult to polarize. Hence, we only considered genes with large PBS_4_ values in TSI, GBR, and FIN populations for these analyses. To globally assess functional modules contributing to population-specific sequence and expression differentiation in these European populations, we utilized annotation data from the Gene Ontology (GO) Consortium (Ashburner et al. 2000; GO Consortium 2018). In particular, GO terms classify genes by their molecular functions, cellular components, and biological processes (Ashburner et al. 2000; GO Consortium 2018). Therefore, to study relationships of population-specific sequence and expression differentiation to gene functions, we ranked genes by their sequence and expression PBS_4_ values in each population, performed GO enrichment analysis on ranked lists, and extracted significantly overrepresented GO terms (Tables S6-S14; see Materials and Methods for details).

First, we examined GO enrichment for sequence PBS_4_ values in TSI, GBR, and FIN populations (Tables S6-S8). Though the three populations do not share any GO terms, they each show enrichment in terms related to neuronal processes. In TSI, the most enriched GO term is “histone methylation”, a process whereby methyl groups are attached to histone proteins. Several other enriched terms are also related to methylation, which is interesting because methylation is often used as a signal for gene activation or silencing (Jones 2012), suggesting that population-specific sequence differentiation is related to epigenetic processes in TSI. Moreover, a few enriched GO terms in TSI are related to dynamics of cell membranes, which are common targets of adaptation (Hamblin and Di Rienzo 2000; Sabeti et al. 2006). In GBR, the most enriched GO term for sequence PBS_4_ values is “branching involved in labyrinthine layer morphogenesis”, a process whereby the branches of fetal placental villi are generated and organized. Therefore, increased population-specific sequence differentiation in GBR may be related to reproduction and fetal development. This is particularly intriguing because, though natural selection frequently targets reproductive phenotypes, often they are male-specific (Pröschel et al. 2006; Ellegren and Parsch 2007; Assis and Bachtrog 2013; Harrison et al. 2015). Yet this GO term specifically relates to female-specific functions, indicating that population-specific adaptation may be associated with female reproductive phenotypes in GBR. In FIN, the most enriched GO term for sequence PBS_4_ values is “ncRNA transcription”, a process whereby noncoding DNA is transcribed into RNA. Though noncoding RNAs are not translated into proteins, they often play important roles in gene regulation and expression (He and Hannon 2004; Mercer et al. 2009). Also interesting is that many other enriched GO terms in FIN are related to reproduction, and specifically to male gamete generation and spermatogenesis, which are often targets of adaptation (Li et al. 2002; Zhou and Bachtrog 2012).

Next, we examined GO enrichment for expression PBS_4_ values in TSI, GBR, and FIN populations (Tables S9-S14). Using *P*_ST_ with *h*^*2*^ = 0.5 and with *h*^*2*^ = 1 yielded similar results, consistent with our expectations based on previous comparisons (see Figures 1 and 2). However, a surprising finding was the abundance of enriched GO terms for expression PBS_4_ values relative to those for sequence PBS_4_ values. Perhaps as a result, many enriched GO terms are shared by the three populations: “cell surface receptor signaling pathway”, “developmental process”, “positive regulation of response to stimulus”, “regulation of cell communication”, “regulation of immune system process”, “regulation of multicellular organismal process”, and “signal transduction”. Though most of these GO terms are quite general and difficult to interpret, it appears that population-specific expression differentiation in Europeans is often related to processes involved in signal transduction, immunity, reproduction, and development. Moreover, many related GO terms are enriched in Europeans, including those with the most enrichment in each population. The abundance of these GO terms is not surprising, particularly as such processes are frequent targets of natural selection (Barreiro and Quintana-Murci 2010; Fumagalli et al. 2011; Enard et al. 2016).

To glean further insight into specific genes potentially driving population-specific sequence and expression differentiation in Europeans, we performed literature searches on genes with the largest sequence and expression PBS_4_ values in each population. The gene with the largest sequence PBS_4_ value in both TSI and GBR is *MCM6*, or Minichromosome Maintenance Complex Component 6. *MCM6* is part of a protein complex essential for the initiation of eukaryotic genome replication (Labib et al. 2000). Moreover, two of its introns contain enhancers of its upstream gene *LCT*, or Lactase, one of which has a mutation that is prevalent in European populations and is thought to confer lactose tolerance in adulthood (Enattah et al. 2002; Troelsen et al. 2003). Consistent with this hypothesis, several genetic studies have identified *MCM6* and *LCT* as targets of recent positive selection in Europeans (Bersaglieri et al. 2004; Voight et al. 2006; Ranciaro et al. 2014; Cheng et al. 2017). Moreover, the gene with the second-largest sequence PBS_4_ value in GBR is *LCT*. In contrast to the other two European populations, the gene with the largest sequence PBS_4_ value in FIN is *NPIPA2*, or Nuclear Pore Complex Interacting Protein Family Member A2. *NPIPA2* is part of the nuclear pore complex of the cell membrane, which regulates exchange between the nucleus and cytoplasm (Strambio-De-Castillia et al. 2010). Polymorphisms in *NPIPA2* are associated with several diverse classes of cancer (Dingerdissen et al. 2017), and its deletion is common in early-onset colorectal cancer (Perea et al. 2017). Moreover, both the localization of *NPIPA2* to the cell membrane and its function in transport across the membrane make it a likely target of natural selection (Tang and Presgraves 2009; Tracy et al. 2010).

The gene with the largest expression PBS_4_ value (for *P*_ST_ with *h*^*2*^ = 0.5 and *h*^*2*^ = 1) in TSI is *PRRX1*, or Paired Related Homeobox 1. *PRRX1* is a DNA-associated protein that functions as a transcription coactivator and is involved in the establishment of diverse mesodermal muscle types during embryonic development (Martin et al. 1995). In particular, *PRRX1* is thought to play a critical role in craniofacial muscle development, as its variants are associated with several diseases that result in facial and neck malformations linked to sleep apnea (Martin et al. 1995; Urbizu et al. 2013). Further, *PRRX1* has also been associated with numerous cancers (Takahashi et al. 2013; Guo et al. 2015; Hirata et al. 2015; Jurecekova et al. 2016; Takano et al. 2016; Zhu et al. 2017), and is thought to mediate metastasis, or the migration and invasion of cancer cells into diverse tissues (Ocaña et al. 2012; Takahashi et al. 2013; Guo et al. 2015; Zhu et al. 2017). In GBR, the gene with the largest expression PBS_4_ value (for *P*_ST_ with *h*^*2*^ = 0.5 and *h*^*2*^ = 1) is *PRKCB*, or Protein Kinase C Beta. *PRKCB* is involved in a diversity of cellular signaling pathway, including B cell activation during immune response (Lutzny et al. 2013), apoptosis (Reyland 2009), and autophagy (Patergnani et al. 2013). As a result, mutations in *PRKCB* are associated with numerous common diseases, including several cancers (Lutzny et al. 2013; Wallace et al. 2014; Antal et al. 2015) and autoimmune diseases (Han et al. 2009a; Sheng et al. 2010; Kawashima et al. 2017). The association with autoimmune diseases is particularly intriguing, as such genes are often identified as targets of recent positive selection (Barreiro and Quintana-Murci 2010; Ramos et al. 2014). It is hypothesized that mutations that cause autoimmune response today may have provided pathogen resistance in the past (Barreiro and Quintana-Murci 2010). Last, in FIN, the gene with the highest expression PBS_4_ value for *P*_ST_ with *h*^*2*^ = 0.5 in FIN is *VDR*, or Vitamin D Receptor, whereas the gene with the highest expression PBS_4_ value for *P*_ST_ with *h*^*2*^ = 1 is *PLAC8*, or Placenta Specific 8. *VDR* interacts with vitamin D in the small intestine to facilitate calcium transportation into circulation (Holick 2006), and has been associated with vitamin D-dependent rickets (Holick 2006; Wagner and Greer 2008) and osteoporosis (Holick 2004). Skin exposure to solar ultraviolet radiation (UVR) produces about 90% of the vitamin D that an individual requires (Holick 2006), and living at high latitudes has been associated with vitamin D deficiency due to decreased UVR (Kimlin 2008; Chaplin and Jablonski 2009). Therefore, it is possible that expression differentiation of *VDR* may contribute to high latitude adaptation in FIN. *PLAC8* is also an interesting candidate, as it was first identified in human dendritic cells (Rissoan et al. 2002) and was later found to be expressed in interstitial extravillous trophoblast cells in the placenta, playing a key role in promoting their invasion and migration (Chang et al. 2018). *PLAC8* also facilitates the epithelial-to-mesenchymal transition (EMT) in colon cancer cells (Li et al. 2014) and has been associated with autoimmune diseases (Orrù et al. 2013). Therefore, expression differentiation of *PLAC8* in FIN may be related to its role in reproduction or disease.

## 3. Discussion

Identifying drivers of human phenotypic variation is crucial to understanding adaptive events that occurred in the past, as well as to developing population- and individual-targeted treatments for diseases in the future (Jorde et al. 2001; Sabeti et al. 2002; Akey et al. 2004). Though previous research (Sabeti et al. 2002; Akey et al. 2004; Voight et al. 2006) has made use of abundant whole-genome and polymorphism data for many human populations (International HapMap 3 Consortium 2010; The 1000 Genome Projects Consortium 2015) to answer this question, simultaneously studying sequence and expression differentiation may provide unique insights into direct phenotypic targets of natural selection. In particular, it is thought that phenotypic evolution more often occurs through changes in gene regulation and expression, rather than their protein-coding sequences (King and Wilson 1975; Wang et al. 1996; Wray et al. 2003; Carroll 2005; Carroll 2008; Raj et al. 2010). For this reason, gene expression differentiation might better reflect phenotypic variation. Therefore, a major advantage of the present study is that we utilized from both sequence and expression data to address questions about population-specific differentiation in humans. Further, results from our combined analysis suggest that population-specific sequence and expression differentiation in humans may be attributed to several important biological processes, most notably immunity and reproduction, and also pinpoint many candidate genes for further study of human phenotypic variation in adaptation and disease.

A potential limitation of our study is the usage of RNA-seq data obtained from lymphoblastoid cell lines. In particular, the enrichment of immune-related functions in genes with high levels of population-specific expression differentiation may be attributed to usage of this cell line, rather than reflecting widespread evolutionary patterns of immunity genes across tissues. Yet it is important to note that associations between increased population-specific expression differentiation and immunity are consistent with previous findings. Specifically, immunity genes are among the fastest evolving genes in the human genome, likely due to adaptations to rapidly changing environments and introductions of novel pathogens (Barreiro and Quintana-Murci 2010; Fumagalli et al. 2011; Enard et al. 2016). Therefore, though observed patterns of population-specific expression differentiation may not be representative of those in other cell types, genes with high population-specific expression differentiation should be further studied to examine their potential roles in human evolutionary history and disease. Regardless, future availability of RNA-seq data for multiple cell or tissue types in several populations will be invaluable for capturing complex patterns of population-specific expression differentiation and pinpointing genic targets of phenotypic variation among human populations.

Another caveat of these RNA-seq data (Lappalainen et al. 2013) is that TSI, GBR, and FIN are closely related European populations. Detection of population-specific differentiation is inherently difficult, as strong positive selection is thought to be rare in recent human evolution (Hernandez et al. 2011; The 1000 Genome Projects Consortium 2015). In addition, we expect sequence and expression differentiation to be correlated among these populations due to their shared ancestry and possible gene flow. Moreover, genetic and phenotypic differences among distantly related populations are better described than those among closely related populations, making it difficult to interpret our findings in the context of human phenotypes. Therefore, future availability of RNA-seq data from additional populations, particularly those that are more distantly related, will be critical to studying population-specific variation and its role in both human evolution and disease.

Despite the limitations of these data, a major advantage of our study is the design of PBS_4_, a novel summary statistic that can be used to estimate population-specific sequence or expression differentiation in four populations. In particular, PBS_4_ requires minimal assumptions about the data and can be used to rapidly estimate population-specific sequence or expression differentiation on a genome-wide scale. Further, because PBS_4_ utilizes data from four populations, branch lengths are more likely to represent true population-specific differentiation than differentiation that occurred ancestral to two populations, as is possible in a three-population scenario (Assis 2018). Therefore, though the dataset used in our study is not ideal in many respects, PBS_4_ can easily be applied to existing or future datasets to quantify population-specific sequence or expression differentiation in humans and other species. In particular, we envision that application of PBS_4_ to future human RNA-seq data from multiple cell lines or tissues and in many populations of varying divergence levels will shed light on complex questions about human evolutionary history and disease processes.

## 4. Materials and Methods

### 4.1 Population-genetic analyses

We downloaded the 1000 Genomes Project phase 3 dataset (The 1000 Genome Projects Consortium 2015) for TSI, GBR, FIN, and YRI populations from ftp://ftp.1000genomes.ebi.ac.uk/vol1/ftp/. After filtering out insertions, deletions, and single-nucleotide polymorphisms with Minor Allele Frequencies (MAF) less than 0.01, we were left with 15,145,123 SNPs. We calculated Hudson’s *F*_ST_ for each SNP (Reynolds et al. 1983; Weir and Cockerham 1984; Bhatia et al. 2013). Then, we combined SNPs within the entire annotated region of each gene and computed the “ratio of averages” for Hudson’s *F*_ST_ (Reynolds et al. 1983; Weir and Cockerham 1984; Bhatia et al. 2013). Because negative *F*_ST_ values are not defined (Wright 1951) and have no biological interpretation (Akey et al. 2002), we followed the standard of setting all negative *F*_ST_ values to zero (e.g., Nei 1990; Akey et al. 2002).

### 4.2 Gene expression analyses

We obtained RNA-seq data from lymphoblastoid cell lines in TSI, GBR, FIN, and YRI populations from the GEUVADIS project (Lappalainen et al. 2013). We excluded data from the population of Utah Residents with Northern and Western European Ancestry (CEU) because they were collected from an older cell line and display expression patterns that are inconsistent with their demographic history (Yuan et al. 2015). We quantified the abundance of transcripts in the remaining 371 individuals (93 in TSI, 94 in GBR, 95 in FIN, and 89 in YRI) using featureCount (Liao et al. 2013) with the long reads option (-L) and the GRCh37 human genome (Zerbino et al. 2017) as our reference. To normalize count data, we used the “median ratio method” method (Anders and Huber 2010) by implementing the estimateSizeFactors function in DESeq2 (Love et al. 2014). Next, we calculated the Fragments Per Kilobase of transcript per Million mapped reads (FPKM) of each gene using DESeq2 (Love et al. 2014). We removed genes that contained fewer than 10 reads in each sample (lowly-expressed), were located on sex chromosomes, or were not protein-coding. For the remaining 13,075 genes, we log-transformed their FPKM values by log(FPKM + 1). We computed the *P*_ST_ for each gene as 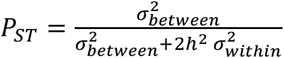 (Leinonen et al. 2006), where 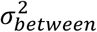 is expression variance between populations, 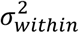 is expression variance within populations, and *h*^*2*^ is heritability. For our analysis, we used *h*^*2*^ = 0.5 and *h*^*2*^ = 1 as was done previously (Leinonen et al. 2006), though we noted that the patterns in Figure 1 do not change as a function of *h*^*2*^. When *h*^*2*^ = 1, *P*_ST_ reduces to *Q*_ST_ (Spitze 1993), another common metric for differentiation of quantitative traits between populations.

### 4.3 Phylogenetic analyses

To infer population trees, we first built gene trees using the NEIGHBOR program in the PHYLIP package (Felsenstein 1993). We constructed gene trees using either *F*_ST_ or *P*_ST_ as input distances between populations. Application of the UPGMA algorithm in the NEIGHBOR program yielded totals of 12,977 gene trees for *F*_ST_ and 13,075 gene trees for *P*_ST_. Next, we used gene trees as input for the CONSENSE program in the PHYLIP package (Felsenstein 1993) and obtained rooted population trees supported by the majority of gene trees based on *F*_ST_ and *P*_ST_. Specifically, the nodes in gene trees are included if they continue to resolve the population tree and do not contradict with more frequently occurring nodes. The number above each node in Figure 1 therefore represents its occurrence in all gene trees.

### 4.4 Calculation of PBS_4_

We first computed the sequence or expression distance between populations as *E*_A,B_ = − log (1 − *Z*_ST_(A,B)), following the approach of Cavalli-Sforza (Cavalli-Sforza 1969), where *Z*_ST_ represents either *F*_ST_ or *P*_ST_ between populations A and B. We used these as input for calculations of sequence and expression PBS_4_ values. Negative branch lengths were set to zero.

### 4.5 GO enrichment analyses

We performed all GO analyses on ranked lists of genes with the GOrilla tool (Eden et al. 2007; Eden et al. 2009). For each run, we output results for all enriched GO categories (process, function, and component) and set the *P*-value threshold to *P* = 10^−3^.

### 4.6 Statistical analyses

All statistical analyses were performed in the R software environment (R Core Team 2013). Two-sample permutation tests were used to assess pairwise differences between all groups compared in Figure 3. For each test, we performed 1,000 permutations, using the difference between medians of groups as the test statistic. In particular, we computed the difference between the medians of the two groups for each permutation, and the *P*-value of the permutation test as the proportion of times the absolute value of this difference was greater than or equal to the absolute value of the observed difference in the data. The significance of correlation coefficients shown in Table S1-2 were assessed via Student’s *t* tests.

## Acknowledgements

This work was supported by the National Science Foundation (DEB-1555981). Portions of this research were conducted with Advanced Cyber Infrastructure computational resources provided by the Institute for CyberScience at Pennsylvania State University (https://ics.psu.edu).

